# Adaptive evolution of animal proteins over development: support for the Darwin selection opportunity hypothesis of Evo-Devo

**DOI:** 10.1101/161711

**Authors:** Jialin Liu, Marc Robinson-Rechavi

**Affiliations:** Department of Ecology and Evolution, University of Lausanne, Switzerland; Swiss Institute of Bioinformatics, Lausanne 1015, Switzerland

## Abstract

A driving hypothesis of Evo-Devo is that animal morphological diversity is shaped both by adaptation and by developmental constraints. Here we have tested Darwin’s “selection opportunity” hypothesis, according to which high evolutionary divergence in late development is due to strong positive selection. We contrasted it to a “developmental constraint” hypothesis, according to which late development is under relaxed negative selection. Indeed, the highest divergence between species, both at the morphological and molecular levels, is observed late in embryogenesis and post-embryonically. To distinguish between adaptation and relaxation hypotheses, we investigated the evidence of positive selection on protein-coding genes in relation to their expression over development, in fly *Drosohila melanogaster*, zebrafish *Danio rerio*, and mouse *Mus musculus*. First, we found that genes specifically expressed in late development have stronger signals of positive selection. Second, over the full transcriptome, genes with evidence for positive selection trend to be expressed in late development. Finally, genes involved in pathways with cumulative evidence of positive selection have higher expression in late development. Overall, there is a consistent signal that positive selection mainly affects genes and pathways expressed in late embryonic development and in adult. Our results imply that the evolution of embryogenesis is mostly conservative, with most adaptive evolution affecting some stages of post-embryonic gene expression, and thus post-embryonic phenotypes. This is consistent with the diversity of environmental challenges to which juveniles and adults are exposed.

## Introduction

There are two main models to explain the relationship of development and evolutionary divergence. The early conservation model suggests that embryonic morphology between different species within the same group progressively diverges across development (Von Baer 1828); such groups are usually understood to by phyla in a modern context. In contrast, the hourglass model proposes that middle development (the morphological ‘phylotypic’ period) has the highest morphological similarity (Duboule 1994; Raff 1996). Based on recent genomic studies, both models have some level of molecular support. Some studies support the early conservation model (Roux and Robinson-Rechavi 2008; Artieri et al. 2009), while most recent ones support the hourglass model (Kalinka et al. 2010; Irie and Kuratani 2011; Levin et al. 2012; Quint et al. 2012; Drost et al. 2015; Hu et al. 2017; Zalts and Yanai 2017). And in fact the two models may not be mutually exclusive (Piasecka et al. 2013; Liu and Robinson-Rechavi 2018).

Both the early conservation and hourglass models predict that late development has high evolutionary divergence. This high divergence of late development has been interpreted as a consequence of relaxed developmental constraints, i.e. weaker negative selection. For example, Garstang (1922) and Riedl (1978) suggested that the development of later stages is dependent on earlier stages, so higher divergence should be found in the later stages of development (cited in Irie and Kuratani 2014). Indeed, many studies have found evidence for relaxed purifying selection in late development (Castillo-Davis and Hartl 2002; Roux and Robinson-Rechavi 2008; Artieri et al. 2009; Kalinka et al. 2010; Liu and Robinson-Rechavi 2018). An alternative explanation, however, known as Darwin’s “selection opportunity” hypothesis (Darwin 1871)(cited in Artieri et al. 2009), proposed that highly divergent late development could also be driven by adaptive evolution (positive selection), at least in part. This could be due to the greater diversity of challenges to which natural selection needs to respond in juvenile and adult life than in early and mid-development. Notably, weaker negative and stronger positive selection are not mutually exclusive. For example, Cai and Petrov (2010) found the accelerated sequence evolution rate of primate lineage specific genes driven by both relaxed purifying selection and enhanced positive selection. Necsulea and Kaessmann (2014) suggested that the high evolution rate of testis transcriptome could be caused by both sex-related positive selection and reduced constraint on transcription.

As far as we know, few studies have tried to distinguish the roles of adaptation vs. relaxation of constraints in late development (Artieri et al. 2009), and no evidence has shown stronger adaptive evolution in late development. Yet there is an intuitive case for adaptation to act on phenotypes established in late development, because they will be present in the juvenile and adult, and interact with a changing environment.

In the case of detecting individual gene adaptation, one of the best established methods is using the ratio ω of non-synonymous (dN) to synonymous (dS) substitutions (Yang and Nielsen 1998; Hurst 2002). Because synonymous changes are assumed to be functionally neutral, ω>1 indicates evidence of positive selection. As adaptive changes probably affect only a few codon sites and at a few phylogenetic lineages, branch-site models allow the ω ratio to vary both among codon sites and among lineages (Yang and Nielsen 2002; Zhang et al. 2005). Polymorphism based methods such as frequency spectrum, linkage disequilibrium and population differentiation can also be used to identify changes due to recent positive selection (Vitti et al. 2013).

Since several genes with slight effect mutations can act together to have a strong effect, adaptive evolution can act on the pathway level as well (Daub et al. 2013; Berg et al. 2014). In the case of polygenic adaptation, a gene set enrichment test has successfully been applied to detect gene sets with polygenic adaptive signals (Daub et al. 2013; Daub et al. 2017). This gene set enrichment analysis allows to detect weak but consistent adaptive signals from whole genome scale, unlike traditional enrichment tests which only consider top scoring genes with an arbitrary significance threshold.

In order to estimate the contribution of positive selection to the evolution of highly divergent late development, we have adopted three approaches. First, we used modularity analysis to obtain distinct sets of genes (modules) which are specifically expressed in different meta developmental stages (Piasecka et al. 2013; Levin et al. 2016), and compared the signal of positive selection across modules. Second, we applied a modified “transcriptome index” (Domazet-Loso and Tautz 2010) to measure evolutionary adaptation on the whole transcriptome level. Finally, we used a gene set enrichment approach to detect polygenic selection on pathways, and studied the expression of these gene sets over development. Each approach was applied to developmental transcriptomes from *D. rerio, M. musculus*, and *D. melanogaster* and to results of the branch-site test for positive selection in lineages leading to these species. All the analyses found a higher rate of adaptation in late and in some stages of post-embryonic development, including adult.

## Results

In order to characterize the signal of positive selection, we used the log-likelihood ratio test statistic (ΔlnL) of H_1_ to H_0_ models with or without positive selection, from the branch-site model (Zhang et al. 2005) as precomputed in Selectome on filtered alignments (Moretti et al. 2014), and as used in Roux et al. (2014) and Daub et al. (2017). Briefly, ΔlnL represents the evidence for positive selection, thus a branch in a gene tree with a higher value indicates higher evidence for positive selection for this gene over this branch.

### Modularity analysis

For the modularity analysis, we focused on different sets of specifically expressed genes (modules) in each developmental period. Our expectation is that genes in each module have specific involvement during embryonic development (Piasecka et al. 2013), so different adaptation rates of these genes should reflect a stage specific impact of natural selection. In addition, since the modules decompose the genes into different meta development stages, they allow to avoid the potential bias caused by imbalanced time points in each meta development stage from our transcriptome datasets; e.g., many more “late development” samples in fly than in the other two species studied. For *D. rerio*, we obtained seven modules from our previous study (Piasecka et al. 2013) (Figure S1). For *M. musculus* and *D. melanogaster*, we identified three and six modules respectively (see Methods; Figure S1).

Because not all genes have any evidence for positive selection, we first compared the proportion of genes either with strong evidence (*q*-value < 0.2) or with weak evidence (no threshold for *q*-value; ΔlnL > 0) of positive selection across modules. For strong evidence, the proportion is not significantly different across modules in *M. musculus* and *D. melanogaster* (Figure S2). In *D. rerio*, however, there is a higher proportion in the juvenile and adult modules. For the weak evidence, *D. melanogaster* has a higher proportion in pupae and adult modules, but there is no significant difference in *D. rerio* and *M. musculus* (Figure S3).

We then compared the values of ΔlnL for genes with weak evidence of positive selection (Figure 1). In order to improve the normality of non-zero ΔlnL, we transformed ΔlnL with fourth root (Hawkins and Wixley 1986; Roux et al. 2014; Daub et al. 2017).

**Figure 1:**
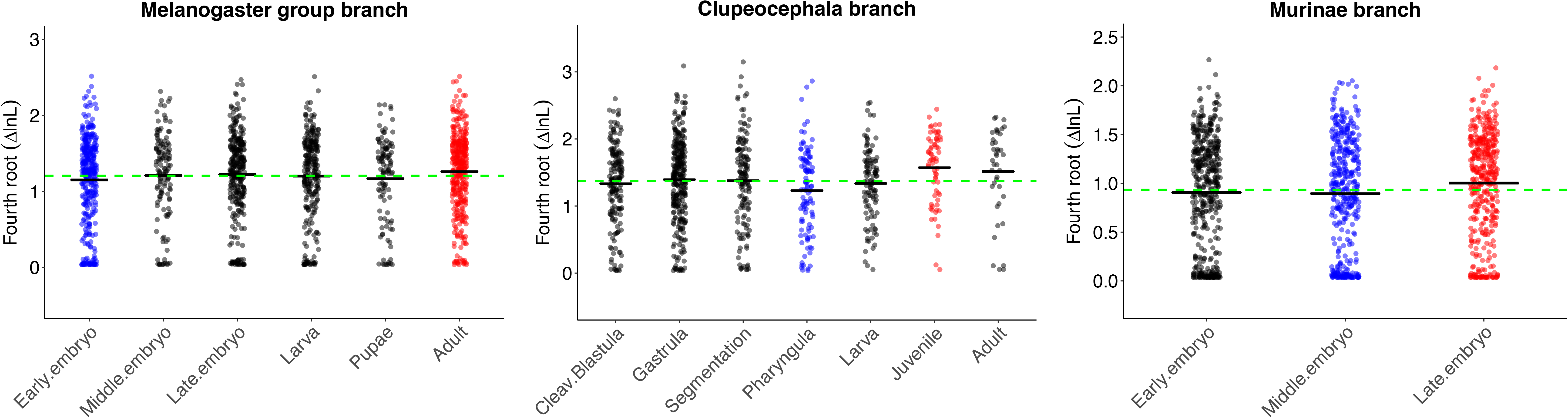
Variation of ΔlnL in different modules. For each module, dots are values of ΔlnL for individual genes and the black line is the mean of ΔlnL. Red (respectively blue) dots indicate modules for which the mean of ΔlnL is significantly (*p*<0.05) higher (respectively lower) than the mean of ΔlnL from all modules. The green dashed line denotes the mean value of ΔlnL from all modular genes.

In *D. rerio*, we detected an hourglass pattern of ΔlnL, at its highest in late modules. Specifically, in the juvenile module, the mean ΔlnL is significantly higher than the mean ΔlnL for all genes (*p*-values reported in Table 1). We note that the adult module also has higher mean ΔlnL, even though it’s not significant. In the pharyngula module, the mean ΔlnL is significantly lower than the mean ΔlnL for all genes, as expected under the hourglass model. In the other modules, the mean ΔlnL is not significantly different from the mean for all genes.

**Table 1.** P-values of randomization test for modular analysis.

|  |  |  |  |  |  |  |  |  |
| --- | --- | --- | --- | --- | --- | --- | --- | --- |
| *D. melanogaster* | Early embryo |  | Middle embryo |  | Late embryo | Larva | Pupae | Adult |
| *p-value* | 0.014 |  | 0.484 |  | 0.263 | 0.435 | 0.213 | 0.018 |
| *D. rerio* | Cleavage/<br>Blastula | Gastrula | Segmentation | Pharyngula |  | Larva | Juvenile | Adult |
| *p-value* | 0.166 | 0.238 | 0.448 | 0.005 |  | 0.273 | 0.003 | 0.066 |
| *M. musculus* | Early embryo |  | Middle embryo |  | Late embryo |  |  |  |
| *p-value* | 0.094 |  | 0.043 |  | 0.001 |  |  |  |

**Table 2.** Candidate pathways enriched with signal of positive selection. We reported all pathways with *g-value* <0.2 after removing overlapping genes (“pruning”) for *D. rerio, D. melanogaster* and *M. musculus*.

| Species | Rank | Pathway | Pathway size<br>before /after<br>pruning | $p$ value<br>before<br>pruning | $q$ value<br>before<br>pruning | $p$ value<br>after<br>pruning | $q$ value<br>after<br>pruning |
| --- | --- | --- | --- | --- | --- | --- | --- |
| *D. rerio* | 1 | Laminin interactions | 12/12 | 2.00E-06 | 1.32E-03 | 2.00E-06 | 0.00E+00 |
|  | 2 | Phenylalanine metabolism | 10/10 | 7.70E-05 | 1.51E-02 | 7.80E-05 | 8.78E-03 |
|  | 3 | Visual phototransduction | 33/33 | 9.10E-05 | 1.51E-02 | 8.30E-05 | 8.78E-03 |
|  | 4 | Metabolism of carbohydrates | 119/118 | 2.02E-04 | 2.23E-02 | 2.37E-04 | 2.97E-02 |
|  | 5 | Gamma carboxylation,<br>hypusine formation and<br>arylsulfatase activation | 18/18 | 1.46E-03 | 8.05E-02 | 1.24E-03 | 1.16E-01 |
|  | 6 | Extracellular matrix<br>organization | 75/61 | 2.00E-05 | 6.62E-03 | 1.50E-03 | 1.16E-01 |
|  | 7 | Acyl chain remodelling of PE | 10/10 | 5.12E-03 | 1.79E-01 | 3.79E-03 | 1.66E-01 |
|  | 8 | Base excision repair | 24/24 | 4.94E-03 | 1.79E-01 | 3.82E-03 | 1.66E-01 |
|  | 9 | Aminoacyl-tRNA biosynthesis | 30/30 | 6.23E-03 | 1.97E-01 | 3.93E-03 | 1.66E-01 |
|  | 10 | Phase II conjugation | 37/30 | 1.08E-03 | 6.92E-02 | 4.00E-03 | 1.66E-01 |
| *D. melanogaster* | 1 | Triglyceride Biosynthesis | 59/59 | 7.60E-05 | 1.57E-02 | 7.60E-05 | 3.32E-02 |
|  | 2 | Glycosaminoglycan<br>degradation | 16/16 | 6.99E-04 | 4.70E-02 | 6.37E-04 | 9.44E-02 |
|  | 3 | Metabolism of porphyrins | 12/12 | 1.57E-03 | 7.31E-02 | 1.47E-03 | 9.62E-02 |
|  | 4 | Detoxification of Reactive<br>Oxygen Species | 17/17 | 1.45E-03 | 7.31E-02 | 1.48E-03 | 9.62E-02 |
|  | 5 | Longevity regulating pathway | 43/28 | 2.83E-02 | 3.12E-01 | 3.37E-03 | 1.54E-01 |
|  | 6 | ECM-receptor interaction | 10/10 | 4.54E-03 | 1.66E-01 | 4.10E-03 | 1.54E-01 |
|  | 7 | Lysine degradation | 25/15 | 1.84E-02 | 2.86E-01 | 5.15E-03 | 1.54E-01 |
|  | 8 | Metabolic pathways | 813/767 | 3.04E-04 | 3.11E-02 | 5.23E-03 | 1.54E-01 |
|  | 9 | Glutathione metabolism | 55/32 | 2.59E-02 | 3.12E-01 | 5.66E-03 | 1.54E-01 |
| *M. musculus* | 1 | Pantothenate and CoA<br>biosynthesis | 16/16 | 9.10E-05 | 7.20E-02 | 9.10E-05 | 5.01E-02 |
|  | 2 | Mineralocorticoid biosynthesis | 10/10 | 1.52E-04 | 7.20E-02 | 1.40E-04 | 5.01E-02 |
|  | 3 | Mitochondrial translation | 72/72 | 2.91E-04 | 7.86E-02 | 2.73E-04 | 5.25E-02 |
|  | 4 | Cytokine-cytokine receptor<br>interaction | 100/100 | 3.41E-04 | 7.86E-02 | 2.78E-04 | 5.25E-02 |

In *M. musculus*, similarly, we found an hourglass pattern of ΔlnL. The late embryo module has a higher mean ΔlnL than all genes, while the middle embryo module has a lower mean ΔlnL than all genes.

In *D. melanogaster*, however, we observed an early conservation pattern of ΔlnL. Specifically, in the early embryo module, the mean ΔlnL is lower than the mean ΔlnL for all genes. In the adult module, the mean ΔlnL is higher than the mean ΔlnL for all genes. There is no significant difference for the other modules.

It should be noted that the patterns reported in this modularity analysis are relatively weak, especially in *D. melanogaster*. After multiple test correction, some of the reported differences are not significant anymore (Table S1).

Overall, these findings suggest that positive selection is stronger on genes expressed in late development or in adult than in early and middle development. It also indicates that ΔlnL on gene modules in different phyla supports different Evo-Devo models (hourglass vs. early conservation).

### Transcriptome index analysis

Although modularity analysis guarantees independence between the sets of genes which are compared, it only considers a subset of genes. This leaves open whether the higher adaptive evolution in late development and adult holds true for the whole transcriptome as well, or just for these modular genes. Additionally, while trends were detected, significance is weak. To consider the composition of the whole transcriptome and to increase our power to detect a signal of positive selection in development, we used a modified “Transcriptome Age Index” (Domazet-Loso and Tautz 2010) to calculate the weighted mean of ΔlnL for the transcriptome. Notably, all expression levels were log-transformed before use, unlike in Domazet-Loso and Tautz (2010). See discussion in Piasecka et al. (2013) and Liu and Robinson-Rechavi (2018), but briefly log-transformation provides insight on the overall transcriptome rather than a small number of highly expressed genes. We named this modified index “Transcriptome Likelihood Index” (TLI). A higher index indicates that the transcriptome has higher expression of transcripts from genes with high ΔlnL between models with and without positive selection.

In *D. rerio*, generally, the pattern resembles an hourglass like pattern (Figure 2). The TLI first decreases and reaches a minimum in the late stage of gastrula (8h), and then progressively increases until adult (ninth month), with finally a slight decline. In addition, in the adult stage, female has higher TLI than male, although the difference is weak. To test whether TLIs are different between developmental periods, we compared the mean TLI of all stages within a period, between each pair of periods (see Methods). We found that middle development has low TLI, early development has medium TLI, late development and maternal stage have very similar high TLI, and adult has the highest TLI. Except late development and maternal stage (*p*=0.24), all pairwise comparisons are significant: *p*<5.7e-07.

**Figure 2:**
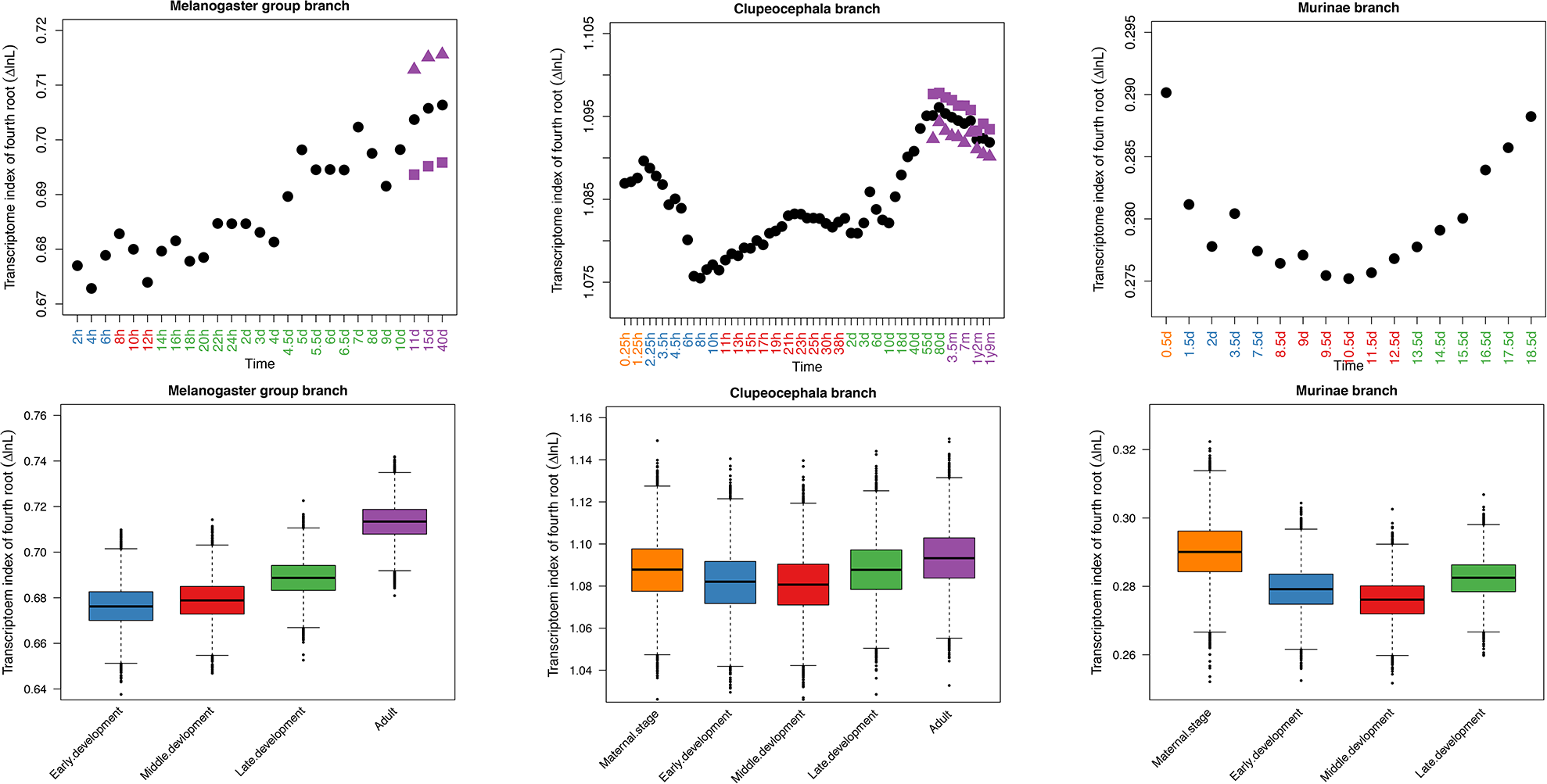
Transcriptome index of ΔlnL (TLI) across development. For sub-figure A, B and C: Orange, blue, red, green and purple time points represent stages within the developmental periods of maternal stage, early development, middle development, late development, and adult, respectively. For the adult stage, the black solid circle represents TLI from average expression between male and female; the purple solid triangle and square represent TLI from only males or females, respectively. For sub-figure D, E and F: Comparison of the TLI (mean TLI of all stages within a period) between any two different periods. Each period has 10,000 pseudo-TLIs which come from random resampling with replacement.

In *M. musculus*, we observed a clear hourglass-like pattern of TLI. For the mean TLI comparison, we found low TLI in middle development, medium TLI in early development, high TLI in late development, and the highest TLI in maternal stage (all pairwise comparisons are significant: *p*<2e-16). Of note, unlike in *D. rerio*, the “late development” here only contains late embryo stages, but no post embryo stages. This may explain why late development has lower TLI than the maternal stage in this dataset.

In *D. melanogaster*, we found the TLI progressively increasing over development, suggesting an early conservation model. Unlike in *D. rerio*, we found that male has higher TLI than female in the adult stage. For the mean TLI comparison, early development has low TLI, middle development has medium TLI, late development has high TLI, and adult has the highest TLI (all pairwise comparisons are significant: *p*<2e-16).

As in the modularity analysis, but with much stronger signal, both *D. rerio* and *M. musculus* support the hourglass model, while *D. melanogaster* follows an early conservation model. Again, from whole transcriptome level, these results indicate that genes with evidence for positive selection are more highly expressed in late development and adult. Interestingly, the maternal stage has a comparable high TLI to late development. This could be related to the maternal stage being dominated by adult transcripts (Tadros and Lipshitz 2009). In this respect (transcriptome evolution), the maternal stage should maybe be regarded as a special adult stage rather than as an early embryonic stage.

### Polygenic selection analysis

Positive selection can be detected at the biological pathway level, even when individual genes within the pathway only fix small effect mutations (Daub et al. 2013; Berg et al. 2014; Daub et al. 2017). Thus, we searched for such signals of positive selection on pathways. Briefly, we calculated the sum of ΔlnL (SUMSTAT statistic) for a pathway, and inferred the significance of this SUMSTAT with an empirical null distribution (Tintle et al. 2009; Daub et al. 2013; Daub et al. 2017). In total, we identified 10, 4 and 9 pathways with a significant signal of positive selection, respectively in lineages leading to *D. rerio, M. musculus* and *D. melanogaster* (*q*-value<0.2, Table2).

The function of these pathways, while not our primary focus, is consistent with adaptive evolution of juvenile or adult phenotypes. First, we found metabolism related pathways in all three species, suggesting pervasive adaptation, possibly related to diet; this is consistent with previous results in primates (Daub et al. 2017). Second, in *D. rerio* and *D. melanogaster*, several pathways are involved in morphogenesis and remodelling of organs (e.g., laminin interactions, extra cellular matrix, ECM-receptor interaction), suggesting potential adaptive evolution of morphological development. Third, there are several pathways involved in aging in *D. melanogaster* and *M. musculus* (e.g., reactive oxygen detoxification, longevity regulation, mitochondrial translation), suggesting potential role of natural selection on modulating lifespan or on metabolic activity. Forth, in *D. rerio*, we detected one pathway related to environmental adaptation: visual phototransduction; adaptations in vision are expected for aquatic species which under a wide variety of visual environments (Sabbah et al. 2010).

If late development and adult are under stronger positive selection at the pathway level as well, we expect genes involved in pathways with a signal of positive selection to be more highly expressed at these periods. Thus we computed the ratio of median expression between positively selected pathway genes and genes included in pathways not positively selected. Since the median expression in the first time point of *M. musculus* is 0, we removed it from our analysis.

In *D. rerio*, the ratio of median expression keeps increasing until the juvenile stage. Then, it slightly decreases (Figure 3). In *M. musculus*, except the first time point, the ratio of median expression also progressively increases. In *D. melanogaster*, there is a small peak in the first time point, but it quickly decreases to minimum within the same developmental period. Then, it keeps increasing until the middle of the larval stage. Finally, for the last development stages, it resembles a wave pattern: decrease, increase and decrease again. Again, we also tested the difference between male and female in adult stages for *D. rerio* and *D. melanogaster*. Unlike the observation in the transcriptome index analysis, here we found that male has higher ratio of median expression than female in both species.

**Figure 3:**
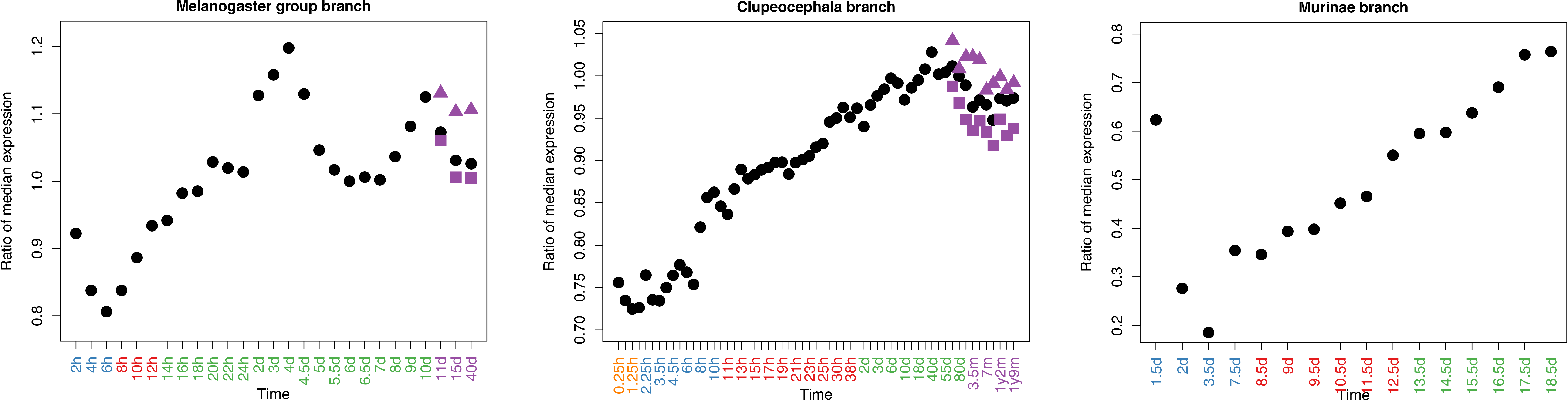
Expression in development for genes involved in pathways enriched with signal of positive selection. Each solid circle represents the ratio of the median expression for genes involved in pathways enriched with signal of positive selection to the median expression for genes involved in pathways without signal of positive selection. Orange, blue, red, green and purple time points represent stages within the developmental periods maternal stage, early development, middle development, late development, and adult, respectively. In adult samples, black solid circles represent ratios generated from average expression of males and females; purple solid triangles and squares represent ratios generated from only males or only females, respectively.

Overall, consistent with previous results, we found that late development and adult tend to express genes involved in pathways enriched for signal of positive selection, indicating that adaptive evolution at the pathway level mainly affects these stages. While there is some signal of early development adaptive evolution on single genes, the later developmental signal is more consistent at the pathway level. Because pathways link genes to phenotypes (Müller 2007; Wray 2007; Tickle and Urrutia 2017), this suggests stronger phenotypic adaptation in late development and adult.

## Discussion

### Correcting confounding factors

Since some non-adaptive factors (such as gene length, tree size (number of branches), and branch length) can be correlated with ΔlnL and affect our results (Daub et al. 2017), we investigated the correlation between ΔlnL and these potential confounding factors. Generally, we found a small correlation between ΔlnL and tree size, but a larger correlation between ΔlnL and gene length or branch length (Figure S4). One explanation for this high correlation between ΔlnL and gene length is that long genes could accumulate more mutations than short genes, so we have more power to detect positive selection with higher number of mutations (Fletcher and Yang 2010; Gharib and Robinson-Rechavi 2013). So, we checked the influence of gene length on our results. Because branch length is inferred from the number of mutations, and higher branch length can be driven by higher evolutionary rate due to positive selection, we did not check further the correlation between ΔlnL and branch length.

In order to investigate whether gene length might have affected our results, for modularity and TLI analysis, we tested whether patterns purely based on gene length are similar to those based on ΔlnL or not. Surprisingly, we found an opposite pattern of gene length, relative to ΔlnL. For modularity analysis, the modules with higher ΔlnL have significantly lower mean gene length than all genes (Figure S5). For transcriptome index analysis, the stages with higher TLI trend to have lower transcriptome index for gene length (Figure S6), suggesting that these stages trend to express shorter genes. These findings imply that the detection of higher positive selection in late development is not driven by gene length.

Immune system genes can bias positive selection analyses, since they evolve under pervasive positive selection (Flajnik and Kasahara 2010). To control for this, we also confirmed our findings after removing immune genes from our analysis (Figure S7).

### Developmental constraint hypothesis and Darwin’s selection opportunity hypothesis

Despite the repeated observation that late development is highly divergent for diverse genomic properties (sequence evolution, duplication, gene age, expression divergence) in diverse animal species (Roux and Robinson-Rechavi 2008; Domazet-Loso and Tautz 2010; Kalinka et al. 2010; Irie and Kuratani 2011; Levin et al. 2012; Piasecka et al. 2013; Drost et al. 2015; Liu and Robinson-Rechavi 2018), the underlying evolutionary forces driving such a pattern remain obscure. The “developmental constraint” hypothesis (Raff 2000; Brakefield 2006) suggests that this high divergence is due to relaxed purifying selection, whereas Darwin’s “selection opportunity” hypothesis proposes stronger positive selection (as discussed in Artieri et al. 2009; Kalinka and Tomancak 2012).

Several studies have found evidence, direct or indirect, to support the importance of developmental constraints (Castillo-Davis and Hartl 2002; Roux and Robinson-Rechavi 2008; Artieri et al. 2009; Kalinka et al. 2010). For example, we (Roux and Robinson-Rechavi 2008) found that genes expressed earlier in development contain a higher proportion of essential genes, and Uchida et al. (2018) found strong embryonic lethality from random mutations in early development. Weaker purifying selection in late development would imply that genes expressed in this period have less fitness impact, which is consistent with the paucity of essential genes. Here and in Liu and Robinson-Rechavi (2017) the branch-site codon model allows us to isolate the contribution of purifying selection to coding sequence evolution. We found indeed that genes under weaker purifying selection on the protein sequence trend to be expressed in late development (Liu and Robinson-Rechavi 2018). This provides direct evidence of relaxed purifying selection in late development.

To the best our knowledge, there has been no direct test of Darwin’s “selection opportunity” hypothesis. One such study, in *D. melanogaster*, was proposed by Artieri et al. (2009). Unfortunately, they only had relatively poor expression data (ESTs) and limited time points (embryonic, larval/pupal and adult), and they did not find any direct evidence of higher positive selection in late development. Since they noticed that the accelerated sequence evolution of genes expressed at adult stage was confounded by male-biased genes, they argued that the rapid evolution observed in late development could be due to specific selective pressures such as sexual selection. A recent study, in *D. melanogaster*, provides indirect evidence: using in situ expression data and population genomic data to map positive selection to different embryonic anatomical structures, Salvador-Martínez et al. (2018) found larva stage enriched with signal of positive selection. Our results clearly provide a quantitative test which supports a role of positive selection in the high divergence of late development. While our sampling is very far from covering the diversity of developmental modes of animals, we show consistent patterns in a placental mammal, a direct development ray-finned fish, and a holometabolous insect. While it is possible that other patterns will be found in species with different development, this shows that adaptation in late development is not limited to one model. We show that this is not due to testis-expressed genes (Figure S8). In addition, in vertebrates, we also found some evidence of adaptive evolution in early development on single genes. This indicates that some changes in early development might be adaptive consequences to diverse ecological niches, as proposed by Kalinka and Tomancak (2012). It should be noted that our results also provide counter evidence to the adaptive penetrance hypothesis, which argues that adaptive evolution mainly occurs in the middle development (Richardson 1999).

### Re-unification of structuralist and functionalist comparative biology

There have been two major approaches to comparative biology since the late 18th century: the structuralist approach (which gave rise to Evo-Devo) emphasizes the role of constraints, and often focuses on investigating spatial and timing variations of conserved structures in distantly related species. In a modern context, the focus is often on comparing developmental genes’ expression between species. The functionalist or adaptationist approach (which gave rise to the Modern Synthesis and most of evolutionary biology) emphasizes the role of natural selection. In a modern context, the focus is often on investigating adaptive mutations. It has been suggested that these two approaches could not be reconciled (Amundson 2007), since the former underscores how mutations generate morphological diversity, while the later underscores whether mutations are fixed by positive selection or not. A good example of the differences between structuralist and adaptationist comes from the debate between Hoekstra and Coyne (2007) and Carroll (2008). As a structuralist, Carroll suggested that mutations affecting morphology largely occur in the *cis*-regulatory regions. However, as adaptationists, Hoekstra and Coyne argued that this statement is at best premature. Their main argument was that they didn’t find that adaptive evolution was more likely occur in *cis*-regulatory elements, but rather in protein coding genes, from both genome-wide surveys and single-locus studies. It is important to note that Carroll’s theory is specific to morphological evolution, but not directly related to evolutionary adaptation. Basically, both sides could be correct, and were mostly discussing different things.

Since both adaptation and structure are part of biology, we should be able to explain both in a consistent manner. Here, we try to bridge positive selection and morphological evolution by combining developmental time-series transcriptomes, positive selection inference on protein coding genes, modularity analysis, transcriptome index analysis, and gene set analysis. From both modularity analysis and transcriptome index analysis, we found that genes highly expressed in late development and adult have higher evidence for positive selection. From polygenic analysis, we found that the expression of positively selected pathways is higher in late development and adult. Overall, these results suggest that higher morphological variation in late development could be at least in part driven by adaptive evolution. In addition, coding sequence evolution might also make a significant contribution to the evolution of morphology, as suggested by Hoekstra and Coyne (2007) and Burga et al. (2017). This is also supported by the observation of tissue-specific positive selection in *D. melanogaster* development (Salvador-Martínez et al. 2018). It should be noted that we do not test here whether regulatory sequence evolution plays a similar or greater role, since we do not have equivalent methods to test for positive selection in regulatory regions.

## Materials and Methods

Data files and analysis scripts are available on our GitHub repository: https://github.com/ljljolinq1010/Adaptive-evolution-in-late-development-and-adult

### Expression data sets

For *D. rerio*, the log-transformed and normalized microarray data was downloaded from our previous study (Piasecka et al. 2013). This data includes 60 stages from egg to adult, which originally comes from Domazet-Loso and Tautz (2010). For *M. musculus*, the processed RNA-seq (normalized but non-transformed) data was retrieved from Hu et al. (2017). This data includes 17 stages from 2cells to E18.5. We further transformed it with log_2_.

For *D. melanogaster*, we obtained processed (normalized but non-transformed) RNA-seq data from http://www.stat.ucla.edu/~jingyi.li/software-and-data.html Li et al. (2014), which originally comes from Graveley et al. (2011). This data has 27 stages from embryo to adult. For the last three stages, since data were available for male and female, we took the mean. We further transformed it with log_2_.

### Branch-site likelihood test data

The log-likelihood ratio (ΔlnL) values of a test for positive selection were retrieved from Selectome (Moretti et al. 2014), a database of positive selection based on the branch-site likelihood test (Zhang et al. 2005). One major advantage of this test is allowing positive selection to vary both among codon sites and among phylogenetic branches. The branch-site test contrasts two hypotheses: the null hypothesis is that no positive selection occurred (H0) in the phylogenetic branch of interest, and the alternative hypothesis is that at least some codons experienced positive selection (H1). The log likelihood ratio statistic (ΔlnL) is computed as 2*(lnLH1-lnLH0). Importantly, in order to mitigate false positives due to poor sequence alignments, Selectome integrates filtering and realignment steps to exclude ambiguously aligned regions.

We used ΔlnL from the Clupeocephala branch, the Murinae branch and the Melanogaster group branch for *D. rerio, M. musculus* and *D. melanogaster* respectively. One gene could have two ΔlnL values in the focal branch because of duplication events. In this case, we keep the value of the branch following the duplication and exclude the value of the branch preceding the duplication.

### Pathways

We downloaded lists of 1,683 *D. rerio* gene sets, 2,269 *M. musculus* gene sets and 1365 *D. melanogaster* gene sets of type “pathway” from the NCBI Biosystems Database (Geer et al. 2009). This is a repository of gene sets collected from manually curated pathway databases, such as BioCyc (Caspi et al. 2014), KEGG (Kanehisa et al. 2014), Reactome (Croft et al. 2014), The National Cancer Institute Pathway Interaction Database (Schaefer et al. 2009) and Wikipathways (Kelder et al. 2012).

### Coding sequence length

We extracted coding sequence (CDS) length from Ensembl version 84 (Yates et al. 2016) using BioMart (Kinsella et al. 2011). For genes with several transcripts, we used the transcript with the maximal CDS length.

### Testis specific genes

Testis specific genes for *M. musculus* and *D. melanogaster* were obtained from a parallel study (Liu and Robinson-Rechavi 2018). The testis specific genes were defined as genes with highest expression in testis and with tissue specificity value ≥ 0.8.

### Immune genes

To control for the impact of immune system genes, we downloaded all genes involved in the “immune response” term (GO:0006955) from AmiGO (Carbon et al. 2009) (accessed on 25.04.2018), and repeated analyses with these genes excluded.

### Phylotypic period

The definition of phylotypic period is based on previous morphological and genomic studies. For *D. melanogaster*, the phylotypic period defined as extended germband stage (Sander 1983; Kalinka et al. 2010); for *D. rerio*, the phylotypic period defined as segmentation and pharyngula satges (Ballard 1981; Wolpert 1991; Slack et al. 1993; Domazet-Loso and Tautz 2010); for *M. musculus*, the phylotypic period defined as Theiler Stage 13 to 20 (Ballard 1981; Wolpert 1991; Slack et al. 1993; Irie and Kuratani 2011).

### Module detection

For D. rerio, we obtained seven modules from our previous study (Piasecka et al. 2013). This is based on the Iterative Signature Algorithm, which identifies modules by an iterative procedure (Bergmann et al. 2003). Specifically, it was initialized with seven artificial expression profiles, similar to presented in Figure S11. Each profile corresponds to one of the zebrafish meta developmental stages. Next, the algorithm will try to find genes with similar expression profiles to these artificial ones through iterations until the processes converges. This method has proven to be very specific, but lacks power with medium or small datasets (<30 time points). For mouse and fly, the sample size is not enough, so we used the method introduced by Levin et al. (2016). Firstly, we generated standardized gene expression for each gene by subtracting its mean (across all stages) and dividing by its standard deviation. Next, we calculated the first two principal components of each gene based on the standardized expression across development. Since the expression was standardized, the genes form a circle with scatter plot (Figure S9). Then, we computed the four-quadrant inverse tangent for each gene based on its principal components, and sort these values to get gene expression order from early to late (Figure S10). Next, we performed Pearson correlation of the standardized expression and idealized expression profile of each module (Figure S11). Finally, for each module, we defined genes with correlation coefficient rank in top 10% as modular genes. Clearly, the genes in earlier modules have higher gene orders (Figure S9).

### Randomization test of modularity analysis

For each module, we randomly choose the same number of ΔlnL from all modular genes (genes attributed to any module in that species) without replacement and calculated the mean value. We repeated this 10000 times and approximated a normal distribution for the mean value of ΔlnL. The *p*-value that the mean value of interested module is higher (or lower) than the mean value from all modular genes is the probability that the randomly sampled mean value of ΔlnL is higher (or lower) than the original mean value of ΔlnL. In the same way, we also estimated the *p*-value of the median ΔlnL value.

### Transcriptome index of log-likelihood ratio (TLI)

The TLI is calculated as:

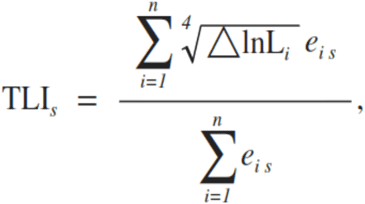

*s* is the developmental stage, ΔlnL_*i*_ is the value of log-likelihood ratio for gene *i, n* the total number of genes and *e_is_* is the log-transformed expression level of gene *i* in developmental stage *s*. Here, we used all ΔlnL values without applying any cut-off on ΔlnL or the associated *p*-value. For genes with ΔlnL<0, we replaced it with 0. For *M. musculus*, we calculated the TLI from a merged data set, instead of computing it on two data sets separately.

### Polynomial regression

For polynomial regression analysis, we keep increasing the degree of polynomial model until no further significant improvement (tested with ANOVA, *p*<0.05 as a significant improvement). For *M. musculus*, since the development time points in transcriptome data set are close to uniformly sampled, we used the natural scale of development time for regression. For *C. elegans, D. melanogaster* and *D. rerio*, however, we used the logarithmic scale, to limit the effect of post-embryonic time points.

### Bootstrap approach for transcriptome index of ΔlnL (TLI) comparison between developmental periods

Firstly, we randomly sampled the same size of genes from original gene set (with replacement) for 10,000 times. In each time, we calculated the TLI of each development stage. Then, we calculated the mean TLI (mean TLI of all stages within a period) for each developmental period (maternal stage, early development, middle development, late development, and adult). Thus, each developmental period contains 10,000 mean TLI. Finally, we performed pairwise Wilcoxon test to test the differences of mean TLI between developmental periods.

### Detection of polygenic selection

We performed a gene set enrichment approach to detect polygenic signals of positive selection on pathways (Ackermann and Strimmer 2009; Daub et al. 2013; Daub et al. 2017). For each pathway, we calculated its SUMSTAT score, which is the sum of ΔlnL of all genes within this pathway. The ΔlnL values were fourth-root transformed. This approach makes the distribution of non-zero ΔlnL approximate normal distribution (Canal 2005; Roux et al. 2014; Daub et al. 2017). So, with fourth-root transformation, we limit the risk that the significant pathways we found be due to a few outlier genes with extremely high ΔlnL. The SUMSTAT score of a pathway is calculated as:

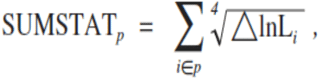

where *p* represents a pathway, and ΔlnL_*i*_ represents the value of log-likelihood ratio for gene *i* within pathway *p*. Pathways less than 10 ΔlnL values were excluded from our analysis. Like in TLI analysis, we used all ΔlnL values and replaced <0 values with 0.

### Empirical null distribution of SUMSTAT

We used a randomization test to infer the significance of the SUSMTAT score of a pathway. To correct for the potential bias caused by gene length, we firstly created bins with genes that have similar length (Figure S12). Secondly, we randomly sampled (without replacement) the same number of genes from each bin, to make the total number of genes equal to the pathway being tested. Thirdly, we computed the SUMSTAT score of the randomly sampled ΔlnL values. We repeated the second and third processes one million times. Fourthly, we approximated a normal distribution for SUMSTAT score of the interested pathway. Finally, the *p*-value was calculated as the probability that the expected SUMSTAT score is higher than the observed SUMSTAT score.

### Removing redundancy in overlapping pathways (“pruning”)

Because some pathways share high ΔlnL value genes, the identified significant pathways might be partially redundant. In other words, shared genes among several pathways can drive all these pathways to score significant. We therefore removed the overlap between pathways with a “pruning” method (Daub et al. 2013; Daub et al. 2017). Firstly, we inferred the p-value of each pathway with the randomization test. Secondly, we removed the genes of the most significant pathway from all the other pathways. Thirdly, we ran the randomization test on these updated gene sets. Finally, we repeated the second and third procedures until no pathways were left to be tested. With this “pruning” method, the randomization tests are not independent and only the high scoring pathways will remain, so we need to estimate the False Discovery Rate (FDR) empirically. To achieve this, we applied the “pruning” method to pathways with permuted ΔlnL scores and repeated it for 300 times. So, for each pathway, we obtained one observed *p*-value (*p*\*) and 300 empirical *p*-values. The FDR was calculated as follow:

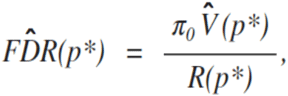

where *π_0_* represents the proportion of true null hypotheses, 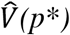 represents the estimated number of rejected true null hypotheses and *R*(*p*\*) represents the total number of rejected hypotheses. For *π*_0_, we conservatively set it equal to 1 as in Daub et al. (2017). For 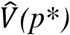, in each permutation analysis, we firstly calculated the proportion of *p*-value (from permutation analysis) ≤ *p*\*. Then, the value of 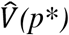 was estimated by the mean proportion of *p*-value (from permutation analysis) ≤ *p*\* for the 300 permutation tests. For *R*(*p*\*), we defined it as the number of *p*-value *(from original analysis)* ≤ *p*\*. For *q*-value, we determined it from the lowest estimated FDR among all *p*-values (from original analysis) ≥ *p*\*.

## Acknowledgements

We thank Sébastien Moretti for help with Selectome data retrieval, the Bgee team for help with expression data retrieval, Barbara Piasecka for help with programming, Josephine T. Daub for help with polygenic selection analysis, and Julien Roux, Elsa Guillot, Iakov Davydov and all members of the Robinson-Rechavi lab for helpful discussions. Part of the computations were performed at the Vital-IT (http://www.vital-it.ch) Center for high-performance computing of the SIB Swiss Institute of Bioinformatics. JL is supported by Swiss National Science Foundation grant 31003A_153341 / 1.

## Author contributions

JL and MRR designed the work. JL performed the data gathering and analysis. JL and MRR interpreted the results. JL wrote the first draft of the paper. JL and MRR finalized the paper.

---

**Table S1.** Multiple test corrected p-values (Benjamini–Hochberg method) of randomization test for modular analysis.

|  |  |  |  |  |  |  |  |  |
| --- | --- | --- | --- | --- | --- | --- | --- | --- |
| *D. melanogaster* | Early embryo |  | Middle embryo |  | Late embryo | Larva | Pupae | Adult |
| *p-value* | 0.056 |  | 0.484 |  | 0.336 | 0.478 | 0.336 | 0.058 |
| *D. rerio* | Cleavage/<br>Blastula | Gastrula | Segmentation | Pharyngula |  | Larva | Juvenile | Adult |
| *p-value* | 0.295 | 0.336 | 0.478 | 0.027 |  | 0.336 | 0.024 | 0.151 |
| *M. musculus* | Early embryo |  | Middle embryo |  | Late embryo |  |  |  |
| *p-value* | 0.188 |  | 0.115 |  | 0.016 |  |  |  |

## Figures legends

**Figure S1: Expression profiles of different modules across development.**

The bold black line represents median expression of modular genes, the two gray lines represent 25th and 75th quantiles of expression of modular genes respectively. The blue, red and green named stages represent early, middle and late modules respectively.

**Figure S2: Proportion of genes with strong evidence of positive selection in each module.**

The number of genes in each module is indicated below each box. The *p*-value from chi-square goodness of fit test is reported in the top-left corner of each graph.

**Figure S3: Proportion of genes with weak evidence of positive selection in each module.**

Legend as in Figure S2.

**Figure S4: Spearman’s correlation between gene properties and ΔlnL.**

Spearman’s correlation coefficient (rho) and adjusted p-value are indicated in the top-right corner of each graph. Loess regression lines are plotted in red.

**Figure S5: Variation of gene length in different modules.**

Legend as in Figure 1.

**Figure S6: Transcriptome index of gene length across development.**

Legend as in Figure 2.

**Figure S7: Transcriptome index of ΔlnL (TLI) for non-immune genes.**

Legend as in Figure 2.

**Figure S8: Transcriptome index of ΔlnL (TLI) for non-testis genes in *M. musculus* and *D. melanogaster*.**

Legend as in Figure 2.

**Figure S9: Scatter plot of genes based on principal component analysis**

Each dot represents one gene, grey dots represent genes not assigned to any modules, blue dots represent genes in early embryo module, red dots represent genes in middle embryo module, blue dots represent genes in late embryo module, pink dots represent genes in larva module, and purple dots represent genes in pupae/adult module. Arrow indicates the gene expression order (from early to late).

**Figure S10: Heat map of gene expression across development**

Genes arranged by the order of expression (from early to late). Generally, the earlier genes have higher expression in earlier stages. Color bar represents the standardized expression value.

**Figure S11: idealized expression profile for each module.**

**Figure S12: Correction of gene length for polygenic selection.**

The red lines indicate the boundaries of bins which contains genes with similar length.

